# MerQuaCo: a computational tool for quality control in image-based spatial transcriptomics

**DOI:** 10.1101/2024.12.04.626766

**Authors:** Naomi Martin, Paul Olsen, Jacob Quon, Jazmin Campos, Nasmil Valera Cuevas, Josh Nagra, Marshall VanNess, Zoe Maltzer, Emily C Gelfand, Alana Oyama, Amanda Gary, Yimin Wang, Angela Alaya, Augustin Ruiz, Cade Reynoldson, Cameron Bielstein, Christina Alice Pom, Cindy Huang, Cliff Slaughterbeck, Elizabeth Liang, Jason Alexander, Jeanelle Ariza, Jocelin Malone, Jose Melchor, Kaity Colbert, Krissy Brouner, Lyudmila Shulga, Melissa Reding, Patrick Latimer, Raymond Sanchez, Stuard Barta, Tom Egdorf, Zachary Madigan, Chelsea M Pagan, Jennie L Close, Brian Long, Michael Kunst, Ed S Lein, Hongkui Zeng, Delissa McMillen, Jack Waters

**Affiliations:** Allen Institute for Brain Science, 615 Westlake Ave N, Seattle WA

**Author notes:** Equal contributions (Naomi Martin, Paul Olsen).

## Abstract

Image-based spatial transcriptomics platforms are powerful tools often used to identify cell populations and describe gene expression in intact tissue. Spatial experiments return large, high-dimensional datasets and several open-source software packages are available to facilitate analysis and visualization. The outputs of spatial transcriptomics platforms are typically imperfect. For example, local variations in transcript detection probability are common. Software tools to characterize imperfections and their impact on downstream analyses are lacking so the data quality is assessed manually, a laborious and often a subjective process. Here we describe imperfections in a dataset of 641 fresh-frozen adult mouse brain sections collected using the Vizgen MERSCOPE. Common imperfections included the local loss of tissue from the section, tissue outside the imaging volume due to detachment from the coverslip, transcripts missing due to dropped images, varying detection probability through space, and differences in transcript detection probability between experiments. We describe the incidence of each imperfection and the likely impact on the accuracy of cell type labels. We develop MerQuaCo, open-source code that detects and quantifies imperfections without user input, facilitating the selection of sections for further analysis with existing packages. Together, our results and MerQuaCo facilitate rigorous, objective assessment of the quality of spatial transcriptomics results.

## INTRODUCTION

The recent advent of spatially resolved molecular imaging methods has enabled the investigation of gene expression patterns within cells in their native tissue context, revealing the organization of transcriptomically-defined cell types (Close *et al*., 2021). Researchers have leveraged emerging spatial technologies to create comprehensive cell-type atlases of a variety of tissue types, including human heart (Asp *et al.,* 2019), breast cancer (Wu *et al.,* 2021), and lung (Madissoon *et al.,* 2023). In brain, in particular, molecular imaging methods have been deployed to unravel the complex spatial relationships of thousands of cell types, resulting in atlases of cell types in adult mouse brain (Zhang *et al.,* 2021; Langlieb *et al*., 2023; Shi *et al*., 2023; Yao *et al.,*2023; Zhang *et al.,* 2023); cell types in adult human brain (Jorstad *et al*, 2023a) and in developing human brain (Braun *et al*., 2023; Velmeshev *et al*., 2023; Kim *et al*., 2023); non-neuronal cells in the mouse nervous system (Zeisel *et al*., 2018); interneurons in mouse, human and non-human primates (Bugeon *et al*., 2022; Chartrand *et al*., 2023; Jorstad *et al*., 2023b; Lee *et al*., 2023); DNA methylation and epigenomics in mouse brain (Liu *et al*., 2023; Zhou *et al*., 2023); and brain cell populations in Alzheimer’s Disease (Gabitto *et al.,* 2023).

Already spatial technologies have enabled many discoveries in biology, but the field of spatial transcriptomics remains immature. Errors may arise during tissue preparation, chemistry, and imaging, resulting in erroneous detection and identification of transcripts. In principle, the sources of these many errors are known. In practice, often it’s unclear how often errors occur, how to best detect and describe the resulting imperfections in the results, and how these imperfections impact downstream analyses such as cell type identification.

For more mature technologies, often the main sources of error are known, there’s consensus on correction strategies, and these corrections are implemented in widely used analysis software suites. In high-throughput RNA sequencing, for example, RNA is fragmented, reverse transcribed to cDNA, and mapped to a known genome, and the number of raw counts per transcript varies with transcript length, GC content, and sequencing depth. No single procedure corrects for all possible errors but various normalization strategies are widely used to minimize within-sample and between-sample effects (Leek *et al*., 2010; Oshlack *et al*., 2010; Conesa *et al*., 2016; Evans *et al*., 2018) and there’s awareness that misinterpretation of results may occur where biological and technical effects are correlated and normalization is inadequate (Conrads *et al*., 2004; Baggerly *et al*., 2004; Liotta *et al*., 2004; Spielman *et al*., 2007; Akey *et al*., 2007, Spielman & Cheung, 2007). Ideally, there would be a consensus around common errors and corrections for spatial transcriptomics, where there is not yet the same emphasis on quality control of results before downstream analyses.

Here we characterize imperfections on MERSCOPE, a commercial platform using Multiplexed Error-Robust single molecule Fluorescence In Situ Hybridization (MERFISH) chemistry (Chen *et al*., 2015; Moffitt & Zhuang, 2016; Moffitt *et al*., 2016). We collected results from 641 adult mouse sections over ∼2 years, developed code to detect and characterize the most common imperfections, and describe the frequency with which each imperfection occurred and its likely impact on cell type identification. Our results indicate that imperfections are common and reduce the accuracy of cell type labels but are rarely severe enough to prevent investigation of the spatial organization of cell populations in adult mouse brain.

Our code, called MerQuaCo, complements existing packages that facilitate the analysis of spatial transcriptomics datasets, including packages focused on data storage and access, e.g. Pysodb, SpatialData (Lin *et al.,* 2024; Marconato *et al.,* 2024); cell segmentation, e.g. cellpose, Baysor (Stringer *et al.,* 2021; Petukhov *et al.,* 2022); and analysis of high-dimensionality spatial data, e.g. Seurat, scanpy, Giotto, squidpy (Sajita *et al*., 2015; Wolf, Angerer & Theis, 2018; Dries *et al.,* 2021; Palla *et al.,* 2022; Hao *et al.,* 2024). Adding MerQuaCo, or comparable procedures, to existing workflows offers an alternative to time-consuming and subjective manual assessment of data quality, streamlining the analysis of large spatial datasets and supporting rigorous, objective assessment of results generated with spatial molecular imaging technologies.

## METHODS

We collected MERFISH results using the Vizgen MERSCOPE platform (https://vizgen.com/products/). Procedures for sample preparation were as described by the Vizgen User Guide (https://vizgen.com/resources/fresh-and-fixed-frozen-tissue-sample-preparation) with modifications in Yao *et al*. (2023). Experiments were conducted on fresh frozen P14-56 mouse brain tissue sectioned at 10 µm onto MERSCOPE coverslips, fixed and permeabilized, hybridized with encoding probes, gel embedded and cleared, and stained with DAPI and polyT to facilitate the identification of somata. Samples were then loaded into the MERSCOPE, which manages sequential fluid exchange and imaging. We excluded 11 experiments with transcripts per µm^2^ per gene <0.0002, yielding 641 sections.

### Pixel Classification

Sample preparation-related errors can arise during MERFISH experiments. Common problems include damage to the section, resulting in the loss of a region of tissue, and detachment from the coverslip, resulting in a localized region of tissue being too far from the coverslip surface to be within the imaging volume.

We built a pixel classifier to quantify the area of each section affected by common prep-related problems. The classifier generates a series of binary masks then combines these masks in a final step, resulting in the classification of each location in the section into one of 5 categories: tissue (tissue within the imaging volume), detachment (tissue present but outside the imaging volume), ventricle (no tissue in the imaging volume but no loss of tissue), damage (no tissue in the imaging volume due to loss of tissue), off-tissue (outside the section).

The intermediate transcript, DAPI, detachment, ventricle, and damage masks were created via mostly binary image operations based on two outputs of the MERSCOPE: the transcript table (which provides the location and gene identity for each transcript) and the DAPI image. To generate these masks we used Random Forest models implemented in ilastik, an interactive image classification, segmentation, and analysis tool (https://www.ilastik.org/).

From the transcript table, we plot an image of transcript counts with 10 x 10 µm pixels, including all transcripts except blanks. The transcript image was converted into a binary mask, the transcript mask, by application of a random forest model trained on binary images from 7 sections, manually annotated to distinguish tissue from all other pixel classes.

From the high resolution DAPI image, we selected plane 0, downsampled in x and y by a factor of 100, and thresholded to remove off-tissue pixels. The resulting modified DAPI image was converted into a binary mask, the DAPI mask, by application of a random forest model trained on modified DAPI images from 6 sections, manually annotated to distinguish DAPI-positive and-negative regions.

The detachment mask was created by subtracting the transcript mask from the DAPI mask. Detached tissue manifests as regions with completely missing transcripts and blurry DAPI signal so the transcript mask excludes detached regions and the DAPI mask includes them.

Our probe panels generally include a few genes expressed preferentially around the boundary of ventricles. We leveraged these genes to distinguish ventricles from regions of tissue damage. We made a list of 11 ventricle-associated genes: Crb2, Glis3, Inhbb, Naaa, Cd24a, Dsg2, Hdc, Shroom3, Vit, Rgs12, Trp73. For each section, we plot transcript density images for all ventricle-associated genes in the probe panel, thresholded each image, and combined all images via an AND operation to create a ventricle outline image (figure 1D). The ventricle outline was summed with the DAPI mask (creating an image with pixel values of 0, 1 and 2) to which we applied a random forest model trained on images from 7 mouse brain sections with annotated ventricles, resulting in the ventricle mask. Of our 641 sections, 20 were imaged with panels lacking any of the 11 ventricle-associated genes. The results in figure 2 were therefore generated from 621 sections.

**Figure 1.**
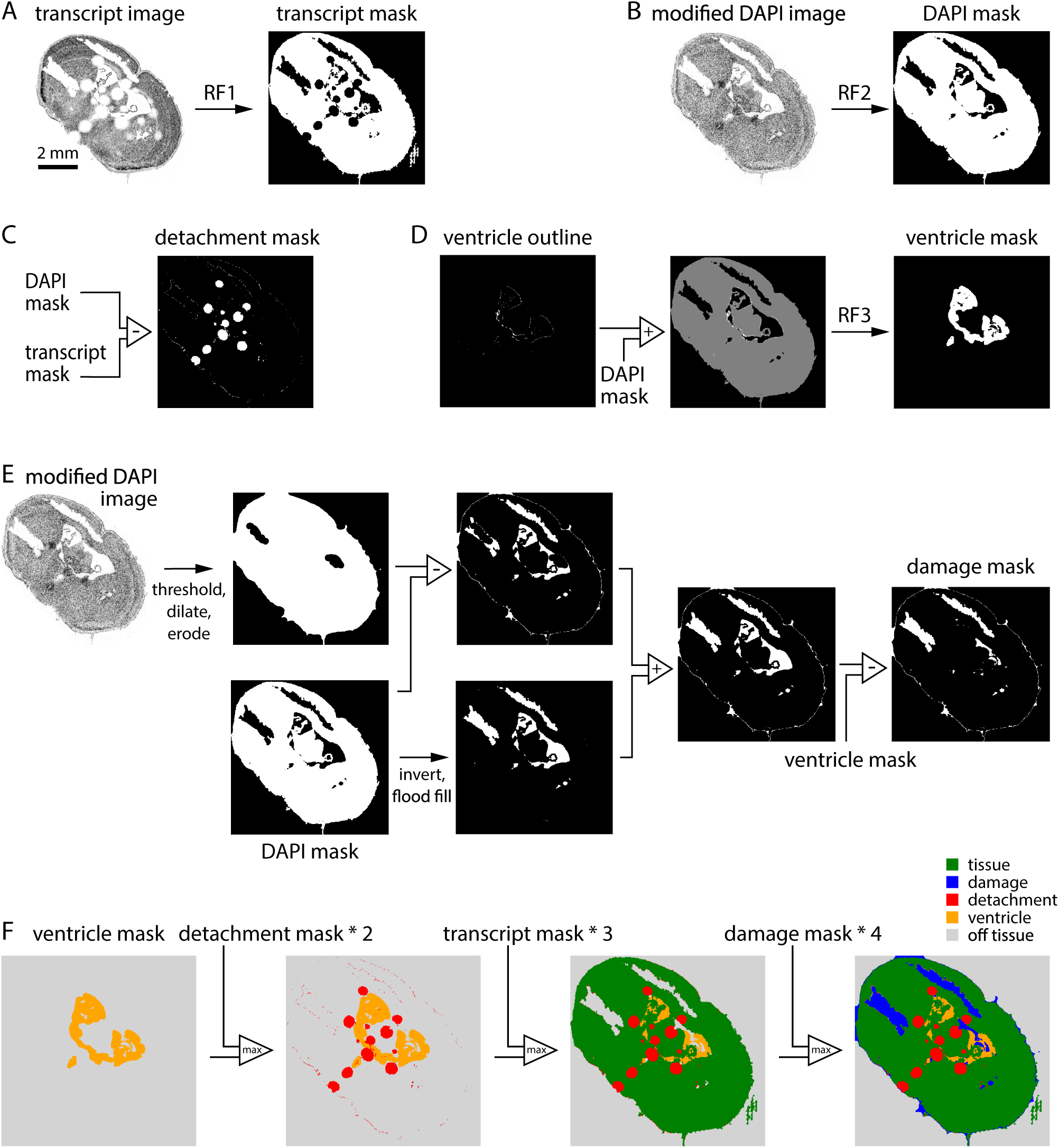
A classifier to assess section integrity. (A) Generation of the transcript mask. The transcript density image was converted to a binary mask using a random forest classifier. (B) Generation of the DAPI mask. The modified DAPI image was converted to a binary mask using a random forest classifier. (C) Generation of the detachment mask. The detachment mask was the difference between DAPI and transcript masks. (D) Generation of the ventricle mask. A binary image summarizing the locations of 11 ventricle boundary genes was summed with the DAPI mask and converted to a binary mask with a random forest classifier. (E) Generation of the damage mask. Two intermediate masks were created via a series of binary operations on the modified DAPI image and DAPI mask, then summed. The ventricle mask was subtracted to remove ventricles. (F) Sequential combination of ventricle, detachment, transcript, and damage masks resulted in the final 5-category image.

**Figure 2.**
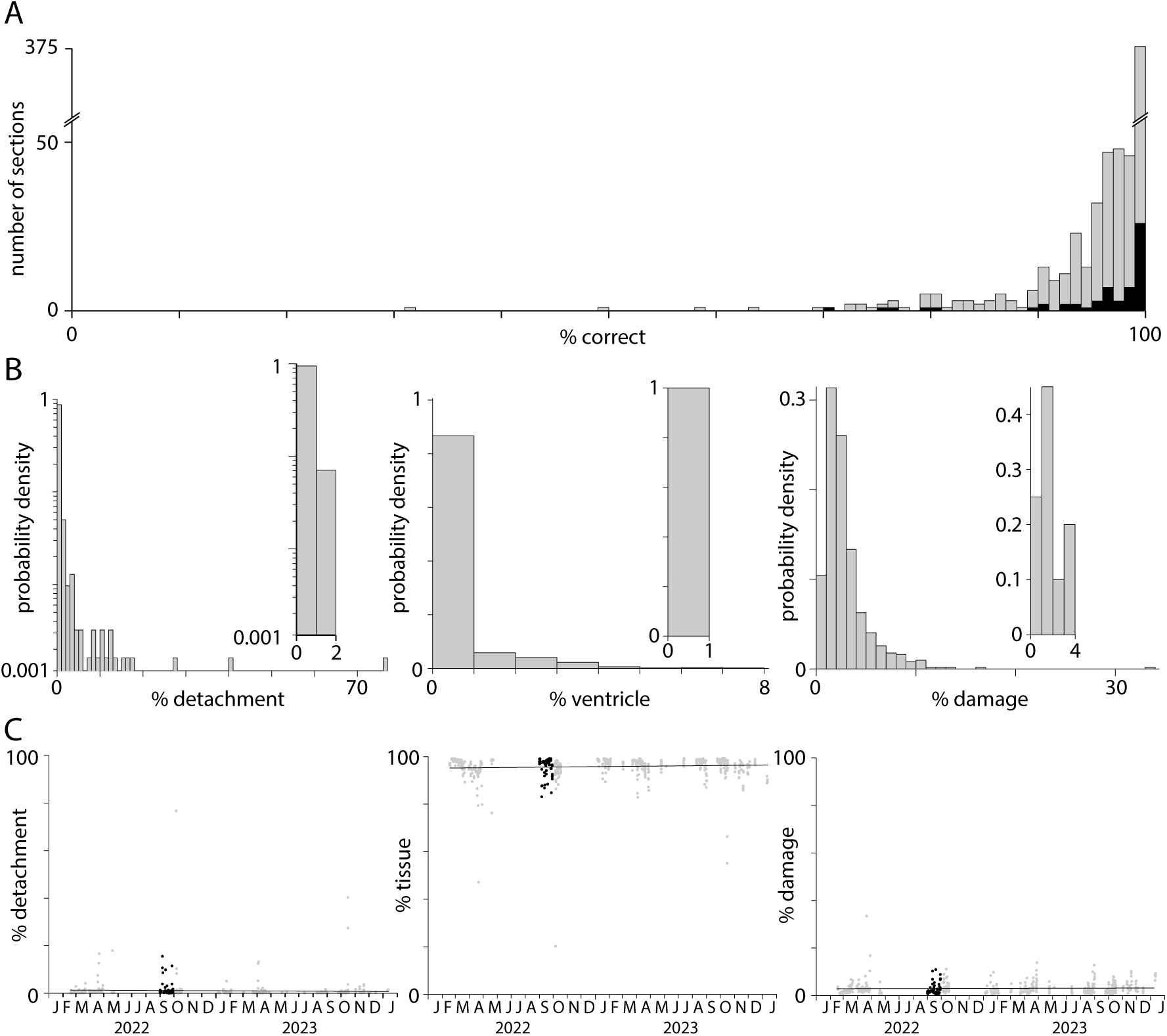
Tissue area, detachment and damage. (A) Accuracy of the pixel classifier, evaluated on a test dataset consisting of a 1 mm^2^ subregion from each of 621 tissue sections (grey). Black: 59 sections in the Yao et al. (2023) Allen Brain Cell Atlas dataset. (B) Probability distributions describing the percentage detachment, ventricle, and damage for each of 621 sections. Insets: false positive distributions calculated for 20 sections without detachment, ventricles, or damage. (C) Percentage of the section identified as on-tissue, detachment, and damage, plot over time (grey), and for the 59 sections in the Yao et al. (2023) Allen Brain Cell Atlas dataset (black).

We investigated two strategies for generating the damage mask. The first began with the modified DAPI image, which was binarized, dilated and eroded, and subtracted from the DAPI mask. This intermediate image often created a mask in which regions of damage were detected but incomplete and with a thin, erroneous strip of damage around the section boundary. The second strategy involved inversion of the DAPI mask followed by elimination of pixels outside the tissue boundary (via a flood fill operation seeded at the origin). This intermediate image often captured ventricles and damage within the section but excluded regions of damage along the section boundary. We summed the two intermediate images, capturing damage within the section and along its boundary. The resulting binary image commonly included ventricles, removed by subtraction of the ventricle mask.

The transcript, detachment, ventricle, and damage masks can contain conflicting pixel labels. To obtain the final pixel classification the 4 masks were combined, with unassigned pixels being off-tissue.

### Perfusion Rate

MERSCOPE outputs a log file that includes the perfusion flow rate (in unspecified units, likely milliliters per minute) at one second intervals during solution exchange. Post-hoc examination of the log file can reveal possible inconsistencies of the flow rate during the experiment, due to blockage of the tubing for example. During a typical MERSCOPE run, median flow was consistently > 1.5 ml/min. Occasionally, solution failed to flow throughout an experiment. We identified experiments with ≥1 solution exchange with a median flow rate <0.5 ml/min. In the 303 experiments for which perfusion log files were available, 30 (10%) had ≥1 compromised reagent solution exchange.

### Data Loss

Like other image-based spatial technologies, MERSCOPE acquires and stitches together many images to map transcript density across a section of tens of millimeters. The absence of an image is readily visualized as a square hole in a plot of transcript locations. To capture this type of data loss, we developed an iterative algorithm that calculates the ratio of transcript counts between every on-tissue field of view (FOV) and its cardinal neighbor FOVs for every gene. In the first iteration, we preliminarily assigned a target FOV as experiencing data loss if the ratio of transcript counts is below 0.15 for any 3 of its 4 cardinal neighbors. We did this to allow the possibility of two neighboring FOVs experiencing data loss; requiring a difference below 0.15 for all 4 cardinal neighbors would result in a high false negative rate in the case of adjacent FOVs with data loss. In a second iteration, we began by removing target FOVs from consideration if the mean transcript counts of their 4 cardinal neighbors is below 100, effectively filtering regions of overall low transcript detection. We then assigned a target FOV as experiencing data loss if the ratio of transcript counts is below 0.15 for all 4 cardinal neighbors, unless one of its neighbors was preliminarily determined to be experiencing data loss in the first iteration, at which point we considered that target FOV to be experiencing data loss in comparison to 3 neighbors.

The third and final iteration aimed to eliminate false positives: FOVs marked as a potential site of data loss after the second iteration where data was not lost. In the third iteration, codebook information was used to determine whether genes in each FOV identified in iteration 2 were in the same round. For each FOV, data loss was excluded where the lost genes were not compatible with the codebook.

### Detection efficiency across the section: periodicity metric

In MERSCOPE images, the transcript count often varies along the x and y axes with a periodicity of ∼200 µm, the size of a field-of-view (FOV). We developed a periodicity metric to describe the uniformity of detection efficiency in the two cardinal (x-and y-) axes across the section. We began by computing a histogram of transcript density in one dimension for one imaging plane (transcripts per µm, either the x-or y-dimension). We divided the histogram into 202 µm segments, approximating the dimensions of a FOV, normalized each segment to its mean transcript density, and calculated the mean of all segments, finally calculating the minimum/maximum density ratio. We repeated this procedure for each of the 7 z-planes and for x-and y-axes, resulting in 14 minimum/maximum ratios. The periodicity metric was the least of these 14 ratios.

### Detection efficiency through the section: p6/p0 ratio

Sections were 10 µm thick. The MERSCOPE images z-planes at 1.5 µm intervals, starting 1.5 µm from the coverslip surface (plane 0). For a 10 µm section, MERSCOPE acquires a stack of seven image planes extending to 10.5 µm (plane 6). Transcript counts often differed across z-planes, generally declining with distance from the coverslip surface. We quantified the gradient by taking the ratio of transcript counts in planes 6 and 0, the p6/p0 ratio. A p6/p0 ratio of 1 corresponds to uniform transcript detection along the z-axis, while a p6/p0 ratio of 0 indicates a failure to detect transcripts in the plane furthest from the coverslip.

### Transcript Density

Transcript density should vary across a tissue section due to differences in gene expression but the mean density per gene should vary little between sections, for sections with comparable RNA quality. The mean transcript density therefore provides an overview of the quality, particularly when benchmarked against a dataset of similar experiments. We calculated transcript density as the mean counts for all on-tissue transcript species (excluding blanks) divided by the area of the on-tissue regions of the sections. Units of transcript density are counts per transcript species per µm^2^.

### Public datasets

We downloaded and analyzed publicly available datasets from four commercial platforms: Vizgen MERSCOPE, 10x Genomics Xenium, NanoString CosMx, and Resolve Molecular Cartography. All datasets include transcript tables, which form the basis for our analyses. All datasets were accessed in July 2024.

The MERSCOPE datasets were animal 1 replicate 2 from the Vizgen MERFISH Mouse Liver Map, a 10 µm thick section imaged with a 347 gene panel (https://info.vizgen.com/mouse-liver-access) and three 10 µm thick coronal sections from three mouse brains in the Vizgen Data Release V1.0 May 2021, imaged with a 483-gene panel (https://info.vizgen.com/mouse-brain-data).

The Xenium datasets were a 5 µm thick formalin-fixed paraffin-embedded (FFPE) coronal mouse brain hemisection from the TgCRND8 mouse model of amyloid precursor protein overexpression, 17.9 months of age, imaged with a 347 gene panel (https://www.10xgenomics.com/datasets/xenium-in-situ-analysis-of-alzheimers-disease-mouse-model-brain-coronal-sections-from-one-hemisphere-over-a-time-course-1-standard) and three 10 µm thick fresh frozen coronal mouse brain sections, imaged with a 248 gene panel (https://www.10xgenomics.com/datasets/fresh-frozen-mouse-brain-replicates-1-standard).

The CosMx dataset is a 10 µm thick FFPE human frontal cortex section imaged with a 6078 gene panel (https://nanostring.com/products/cosmx-spatial-molecular-imager/ffpe-dataset/human-frontal-cortex-ffpe-dataset/).

The Molecular Cartography dataset is a coronal mouse brain hemisection imaged with a 100 gene panel (thickness not stated, https://resolvebiosciences.com/open-dataset/?dataset=mouse-brain-2021)

### Variability of transcript density across sections

In figure 8D, we estimated the variability in transcript counts or density across experiments for MERSCOPE, Xenium and CosMx. Molecular Cartography was excluded since results were available for only one tissue section. Datasets varied in size (2 experiments each in Xenium and CosMx from figure 2A of Cook *et al*. (2023); 3 experiments in Xenium fresh-frozen-mouse-brain-replicates-1-standard dataset; 59 experiments in Yao *et al*. (2023) MERSCOPE dataset) and metric measured (median transcript count per cell in Cook *et al*. (2023); transcript density per gene in Xenium fresh-frozen-mouse-brain-replicates-1-standard dataset; transcript density per gene in Yao *et al*. (2023) MERSCOPE dataset). To enable comparison across metrics and datasets, we calculated the mean coefficient of variation (CV) of all pairwise combinations of experiments. Importantly, the mean CV is independent of sample size and is identical when calculated from transcript density or counts per cell. Equal transcript counts between experiments would result in a CV of 0. CV increases linearly with differences in transcript counts.

### Availability of MerQuaCo

MerQuaCo (for MERSCOPE Quality Control) is a Python package available on Github: https://github.com/AllenInstitute/merquaco.

Documentation: https://merquaco.readthedocs.io/en/latest.

## RESULTS

Our aim was to develop code and characterize data quality for each tissue section processed on our MERSCOPE platforms. We developed code to quantify commonplace imperfections and assess quality by comparing each section to the distribution across 641 mouse brain sections. Our dataset, collected over 2 years on 8 MERSCOPE systems, includes the 59 adult mouse brain coronal sections published in Yao *et al*. (2023) and freely available through the Allen Brain Cell Atlas (https://portal.brain-map.org/atlases-and-data/bkp/abc-atlas). Below, for each imperfection we provide an example, describe our code, and consider the likely effects of each imperfection on cell type identification.

### Tissue preparation

A MERSCOPE experiment starts with sectioning, each tissue section being placed onto a coverslip. After several benchtop chemistry steps, the coverslip is assembled into a flow chamber and then loaded into a MERSCOPE for automated imaging. Common failures during tissue preparation include localized damage resulting in the loss of part of the tissue section, and localized detachment of part of the section from the coverslip. Both result in data loss, the former because tissue is lost and the latter because tissue is present but outside the volume imaged by the MERSCOPE, which extends 10.5 µm from the coverslip surface.

Even in the absence of damage and detachment, some regions of the coverslip lack tissue. These include regions outside the section boundary and tissue-free regions within the section. In brain sections, the latter includes ventricles. Some of the quality metrics we measure in our MERSCOPE experiments, such as transcript density, require that we distinguish regions of the coverslip with and without tissue and we therefore begin our analysis with code that locates tissue.

Ideally, the transcript table would include transcripts only where there’s tissue; there would be no transcripts in regions of the coverslip without tissue. In practice, every experiment includes transcripts where there’s no tissue. Often, the transcript density in some off-tissue regions exceeds that in some on-tissue regions, preventing the use of a simple threshold to identify regions of the coverslip containing tissue. We therefore developed a pixel classifier, which converts the transcript table into a transcript density image and applies a random forest classifier, trained using 10 manually annotated images. The result is the transcript mask, a binary mask which initially classifies each pixel as on-or off-tissue (figure 1A).

We developed our classifier to further categorize off-tissue pixels, resulting in a classification of each pixel into one of five categories: tissue, detached, ventricle, damage, and off-tissue. As inputs, our pixel classifier takes two outputs of the MERSCOPE experiment: the DAPI image and the transcript table. Our strategy was to generate four image masks, each a binary map of one class of pixel (transcript mask, damage mask, detachment mask, ventricle mask) and combine the masks into a single image with our five pixel classes.

Where tissue becomes detached from the coverslip, and is outside the imaging planes of the MERSCOPE, the transcript count is low. Although slightly out of focus, DAPI fluorescence is usually present in regions of detachment (figure 1B, DAPI image). By subtracting the transcript mask from a DAPI mask (generated from the DAPI image with the use of a random forest classifier) we created a detachment mask (figure 1C). For the ventricle mask, we mapped transcripts associated with endothelial cells, which line the ventricle, again using a random forest classifier to convert transcript density to a mask (figure 1D). The damage mask was generated from the DAPI image via a series of binary operations (figure 1E). The final classification was created by summing damage, transcript, detachment, and ventricle masks in sequence (figure 1F).

To assess the accuracy of the pixel classifier, a test dataset of a 1 x 1 mm subregion from each of 621 sections was manually annotated for damage, tissue, detachment, ventricles, and off-tissue, to which the pixel classification results were compared. (The remaining 20 sections were imaged without probes for endothelial cell marker genes.) Pixel classification was >90% accurate for 567 (91%) of 621 subregions (Figure 2A). Typically, the tissue classifier reported <10% detachment, <5% ventricles, and <10% damage (figure 2B, 12 sections with detachment >10%, 4 sections with ventricles >5%, 11 sections with damage >10%). The classifier was prone to detect minor detachment, ventricles and damage in their absence. To quantify the false positive rate for detachment, ventricles and damage, we ran the classifier on 20 sections with no detachment, 20 without ventricles, and 20 undamaged sections. False positive rates were <2% detachment, <1% ventricles, and <4% damage (figure 2B, insets). Only 78 (12.6%) of 621 sections had >2% detachment and likely were partially detached from the coverslip during imaging. 117 (18.8%) of 621 sections had >4% damage and likely suffered some tissue loss due to damage during preparation. There was no significant change in detachment, tissue area or damage over ∼2 years of MERSCOPE experiments so a few percent detachment and damage is routine in our MERSCOPE experiments (figure 2C; Pearson correlation coefficients and p-values: detachment-0.043, 0.28; tissue 0.031, 0.45; damage 0.072, 0.073).

In summary, the classifier estimated tissue area, detachment and damage with reasonable accuracy. We used tissue area in the calculation of subsequent metrics, such as transcript density, and the incidence of common problems in tissue preparation to monitor our tissue preparation and handling procedures.

### Differences in transcript density between MERSCOPE experiments

Once imaging is complete the MERSCOPE runs automated image analysis procedures, returning a transcript table, with the locations and gene identity for each transcript, and a cell-by-gene table, with cell locations and a list of transcripts within each soma. Our analyses of data quality focus on the transcript table. Ideally, the probability of detection of an RNA molecule would be invariant: detection probability would be identical in every experiment, through space within each section, and for different genes. Gene-specific differences in detection probability are almost inevitable with probe-based methods in which the number of target sequences and probe binding differs between genes, but we find that detection probability also varies from experiment to experiment, and often through space within each experiment.

Transcript counts often varied substantially between sections, even for two neighboring sections from the same mouse (figure 3A). For closely spaced sections of similar area, probed with the same gene panel, we expect modest inter-section differences in transcript density due to differing expression patterns through the brain. In practice, inter-section differences in transcript density were approximately 2-fold for a series of sections from a single mouse (collected in a single sectioning session and processed over several weeks; figure 3B). Across 641 sections, transcript density was distributed approximately normally with a mean of 0.0056 and standard deviation of 0.0023 transcripts per gene per square micrometer of tissue (figure 3C). Transcript density differed across probe panels but the variability in transcript density changed little through time (figure 3D).

**Figure 3.**
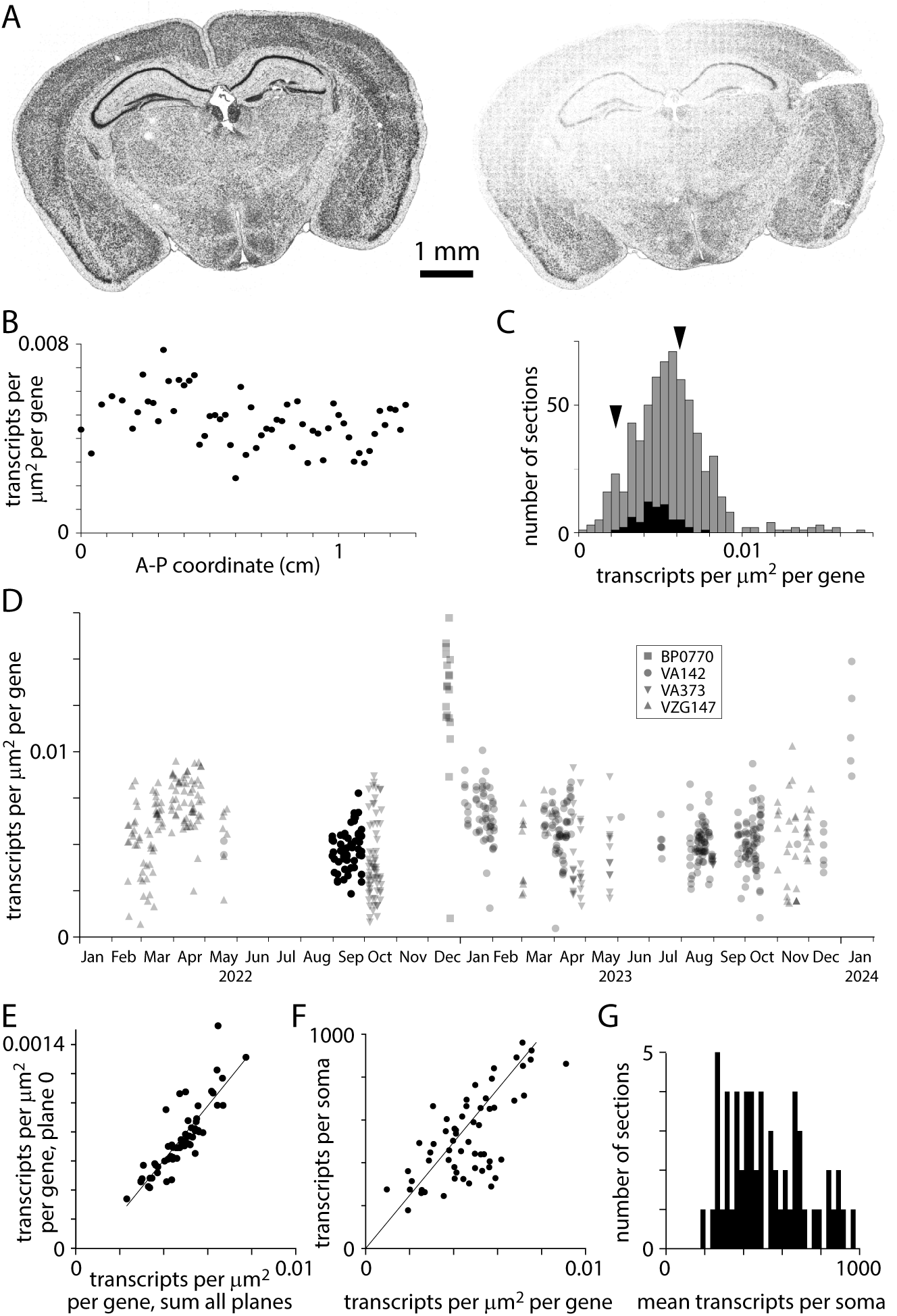
Transcript density. (A) Transcript locations for two neighboring sections from the same mouse brain, separated along the A-P axis by 200 µm. (B) Transcript density across A-P locations for a single mouse. 59 sections in the Yao et al. (2023) Allen Brain Cell Atlas dataset. (C) Histograms of transcript density per transcript species per square micrometer for 641 sections. Summed results from 4 gene panels (VA142, VA373, BP0770, VZG147). Black: 59 sections in the Yao et al. (2023) Allen Brain Cell Atlas dataset (using VA142). Arrowheads: transcript densities for the two sections in A (panel VA142). (D) Transcript density over time. Symbols indicate gene panels. (E) Comparison of transcript densities in plane 0 and summed across all planes. Each data point represents one section from the Yao et al. (2023) dataset. Pearson correlation coefficient 0.86, p = 3.1 x 10^-18^. (F) Mean transcripts per soma vs transcript density. One data point per section, Yao et al. (2023) dataset. Pearson correlation coefficient 0.69, p = 1.07 x 10^-9^. (G) Distribution of transcripts per soma. Yao et al. (2023) dataset.

Why does transcript density vary across sections? Variability was substantial across sections from a single mouse, where tissue quality would have been comparable for all sections, so differences in preparation are unlikely to be responsible. Transcript count in plane 0, just 1.5 µm from the coverslip surface, correlated tightly with total transcript count (figure 3E) and mean transcripts per soma correlated with transcript density (figure 3F) so varying thickness across sections and variable loss of transcripts from the tissue surface during MERSCOPE chemistry are unlikely mechanisms. We conclude that transcript detection efficiency varies across sections, resulting in a broad range of 200-1000 transcripts per soma in a single adult mouse (figure 3G). The mechanism underlying these batch effects remains unclear, but variability in MERSCOPE chemistry, imaging and image analysis all remain candidates.

### Data loss

Transcript density should vary across each section due to differences in gene expression through the tissue. Additionally, artifactual gradients and abrupt changes in transcript density can be introduced by the MERSCOPE. The most abrupt changes in transcript density result from simple data loss. The field of view (FOV) of the MERSCOPE is ∼200 µm x ∼200 µm so to image a tissue section the MERSCOPE tiles many images. If an image is lost, the likely result is the loss of transcripts, readily visualized as a square hole in a plot of transcript locations (figure 4A). The MERSCOPE images three spectrally distinct readout bits in each imaging round. Loss of one of the three images would result in the loss of information on one readout probe. Each readout probe binds to transcripts from tens to hundreds of genes (for panels used here, 60-104 genes). The loss of one bit from the barcode may complicate decoding and decrease the accuracy of detection for many transcript species, but the effect will likely be greatest for transcript species to which the missing readout probe binds. Hence data loss tends to occur for groups of genes, linked by a shared readout probe. Whether the data loss is visible for each of the genes depends on the density of transcripts for each gene in the surrounding regions. In short, data loss typically occurs for multiple but rarely all transcript species and the number of species may not be readily apparent from the transcript table.

**Figure 4.**
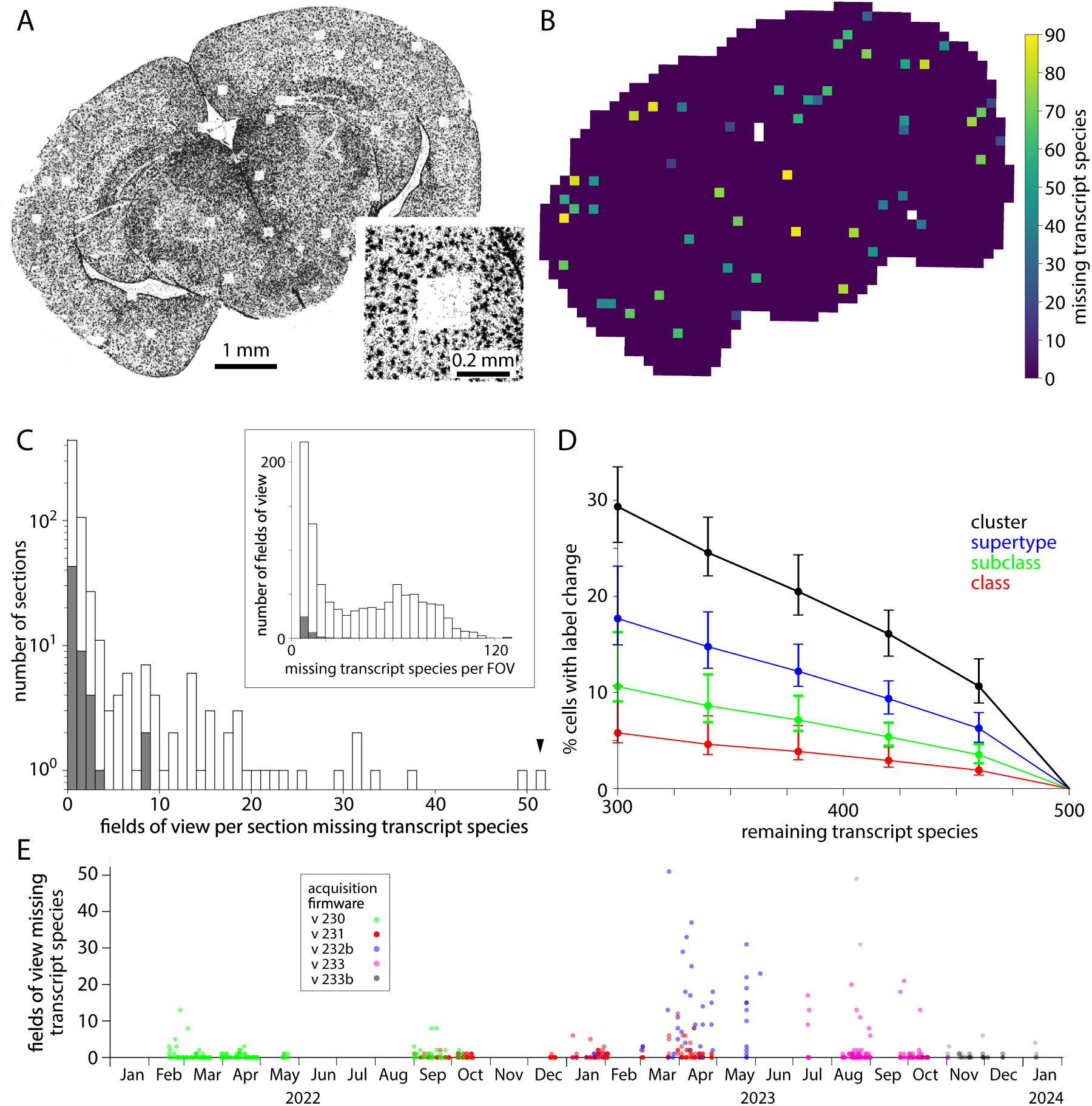
Dropped images cause local data loss. (A) Transcript locations for one transcript species: Gja1. Each point represents one Gja1 transcript. Inset: transcripts around one region of data loss. (B) Output of the data loss detection routine, showing the number of transcript species missing from each FOV. Not all FOVs with dropped genes are missing Gja1. White: off-tissue, as determined by the classifier. (C) Histogram of missing fields of view across 641 mouse sections (grey) and for the 59 sections in the Yao et al. (2023) Allen Brain Cell Atlas dataset (blue). Arrowhead: results for the section in panels A and B. Inset: Number of transcript species missing per affected field of view. (D) Effect of missing transcript species on label transfer. Change in class, subclass, supertype and cluster labels calculated for 10,000 cells from the Yao et al. (2023) Allen Brain Cell Atlas dataset. Median, minimum and maximum % change from 100 trials. Genes to be removed were selected at random. (E) Number of missing fields of view over time and with different acquisition software versions. Each point represents a tissue section.

For each transcript species, we quantified data loss by comparing transcript counts in each field of view to its four cardinal neighbors. A transcript species was considered lost from a field of view if the counts were less than 15% of its cardinal neighbors, with a subsequent false positive correction step. Where a gene was lost its transcript count was 5.1 ± 3.9% of the mean of its cardinal neighbors. As expected, where data was lost from a field of view, often tens of transcript species were missing (figure 4B). Data loss occurred in 201 (31%) of 641 sections and was mostly limited to a few isolated locations with <3 missing fields of view in 133 (66%) of 201 sections (figure 4C). Occasionally more substantial data loss was observed, including loss from up to 51 fields of view for a single section, and 120 transcript species in a single field of view. For sections in the Yao *et al*. (2023) Allen Brain Cell Atlas dataset, 16 of 59 (27%) sections suffered data loss but for no section was there loss from >8 fields of view, with ≤32 transcript species lost from each field of view.

The loss of transcript species reduces the accuracy of cell type labels. For the Yao *et al*. (2023) Allen Brain Cell Atlas dataset with a 500-probe panel (VA142) we calculated the effect of omitting 40-200 transcript species (figure 4D). The loss of 40 transcript species changed the cluster labels of ∼10% of cells so we expect label transfer to be less accurate in fields of view where even one readout bit is lost from the barcode through the loss of an image.

The prevalence of data loss changed substantially over ∼2 years on our MERSCOPEs, as acquisition firmware was updated (figure 4E). Data loss was relatively common with versions 232b and 233 of the acquisition firmware, available in mid-to-late 2023. Data loss has been less common with more recent firmware, such as version 233b, but occurs with all versions of the acquisition firmware.

### Spatially heterogeneity of transcript detection

Ideally, detection efficiency would be uniform throughout the tissue. In practice, detection efficiency is not spatially uniform and there are inter-experiment differences in the non-uniformity. We characterized transcript density in all three cardinal optical axes of the MERSCOPE.

Across the tissue section (in the x and y axes) we expect transcript counts to vary due to differences in gene expression. We observed an additional source of variation: transcript counts varied along x and y axes with a periodicity of ∼200 µm, indicating that detection efficiency varied systematically across each field of view (figure 5A). We characterized the uniformity of detection efficiency with a periodicity metric. We calculated the variation in transcript density across the mean field of view, in x-and y-axes independently, and calculated the minimum/maximum density ratio for x-and y-dimensions for each of the seven z-planes (figure 5B), using the minimum of these 14 values to describe non-uniformity of detection efficiency for each section. A periodicity metric of 1 indicates that detection efficiency was uniform; a value of 0 indicates that no transcripts are detected in part of the field of view.

**Figure 5.**
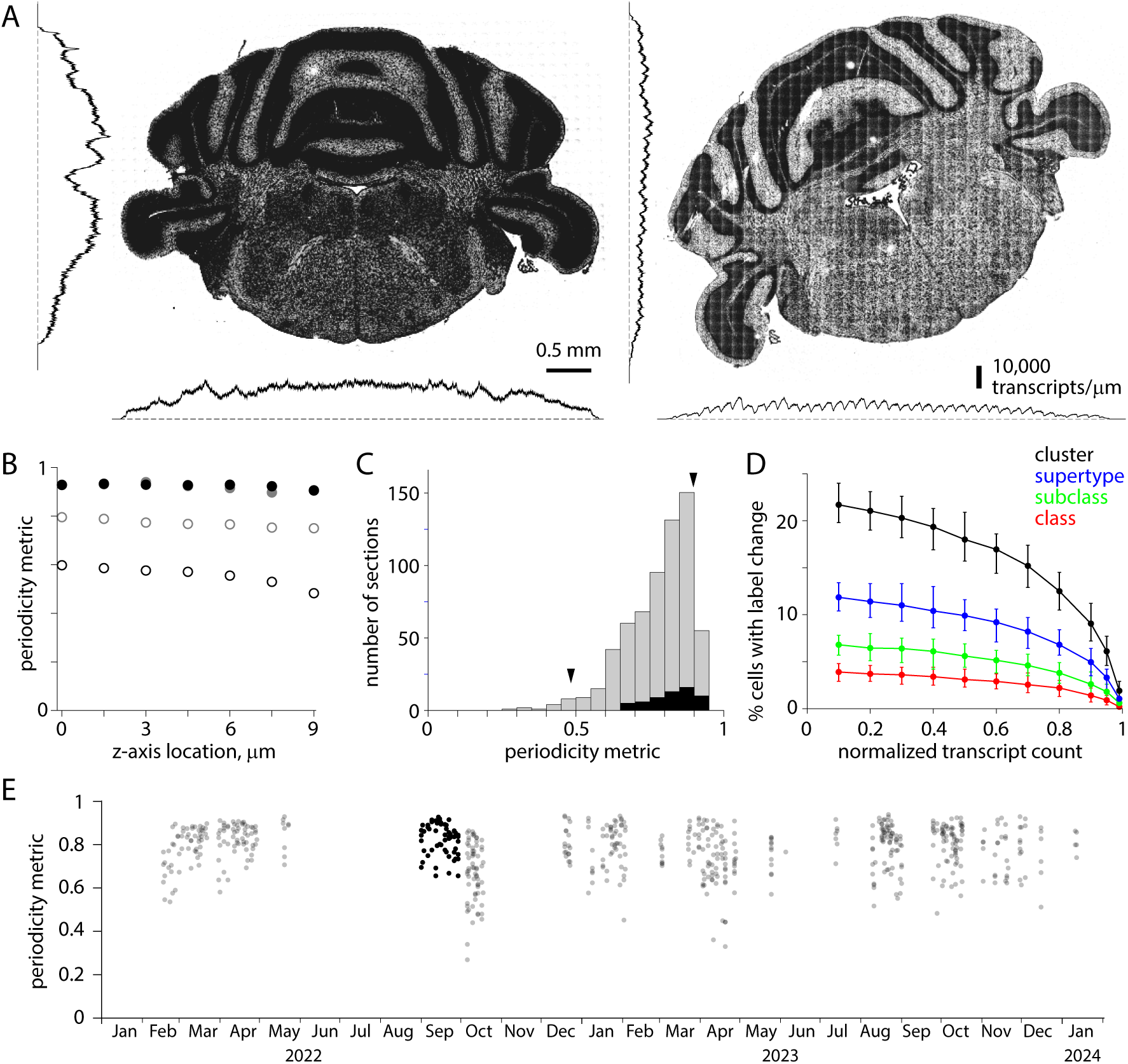
Uneven detection efficiency across each field of view. (A) Transcript locations for two coronal sections from the same brain, separated by 100 µm. To the left and below, transcript densities summed along x and y axes. For the section on the left, changes in transcript density occur at anatomical boundaries with little indication of variations in detection efficiency along x or y axes. For the section on the right, superimposed on differences in genes expression are variations in detection efficiency with a periodicity of 200 µm. (B) Periodicity metric, calculated for each z-plane along x and y axes, for the two sections in A. Filled symbols, left example in A. Open symbols, right example in A. Black and grey, metric along x and y axes, respectively. (C) Histogram of minimum periodicity metrics for 641 sections (grey) and for the 59 sections in the Yao et al. (2023) Allen Brain Cell Atlas dataset (black). Arrowheads, the two sections in A. (D) Effect of reduced detection efficiency on label transfer. Change in class, subclass, supertype and cluster labels for 10,000 cells from the Yao et al. (2023) Allen Brain Cell Atlas dataset (VA142 500 probe panel). Median, minimum and maximum % change from 100 trials. (E) Periodicity metric over time. Black: periodicity metric for the 59 sections in the Yao et al. (2023) Allen Brain Cell Atlas dataset.

Detection efficiency varied across the field of view for all sections, with the variation differing substantially across sections. The median periodicity metric for all 641 sections was 0.80, with a long tail extending towards zero (figure 5C). 40 exhibited a periodicity metric of <0.6. For the 59 sections in the Yao *et al*. (2023) Allen Brain Cell Atlas dataset, the median periodicity metric was 0.84 and the range 0.66-0.93. Based on simulations with the Yao *et al*. (2023) Allen Brain Cell Atlas dataset, we expect reduced detection efficiency, effectively the loss of transcripts, to reduce the accuracy of label transfer. The loss of 20% of transcripts changes the cluster labels of 10-15% of cells, and the loss of 40% of transcripts changes the cluster labels of 15-20% of cells (figure 5D). The differences in detection efficiency across the field of view have changed little over ∼2 years (figure 5E).

Along the optical axis (z axis, perpendicular to the plane of the tissue section), the MERSCOPE acquires images in 7 locations separated by 1.5 µm. Ideally, transcript detection efficiency would be equal in all 7 images, but we routinely observed more transcripts in imaging planes closer to the coverslip than in planes near the tissue-solution interface. Figure 6A shows transcripts from two neighboring sections from the same mouse brain (collected from A-P locations 200 µm apart). In the first section, the transcript count is uniform along the optical axis (figure 6A, B). In the second section, the transcript count is comparable to that in the first section near the coverslip, consistent with similar gene expression in two closely spaced sections, but transcript count declines with distance from the coverslip, to ∼10% 10.5 µm from the coverslip (Figure 6A, B). We quantify homogeneity of detection efficiency along the optical axis with the ratio of transcript counts in planes 6 and 0 (10.5 and 1.5 µm from the coverslip, p6/p0 ratio). Uniform detection efficiency along the optical axis corresponds to a p6/p0 ratio of 1. A p6/p0 ratio of 0 indicates a steep decline in transcript detection with distance from the coverslip, such that no transcripts are detected 10.5 µm from the coverslip.

**Figure 6.**
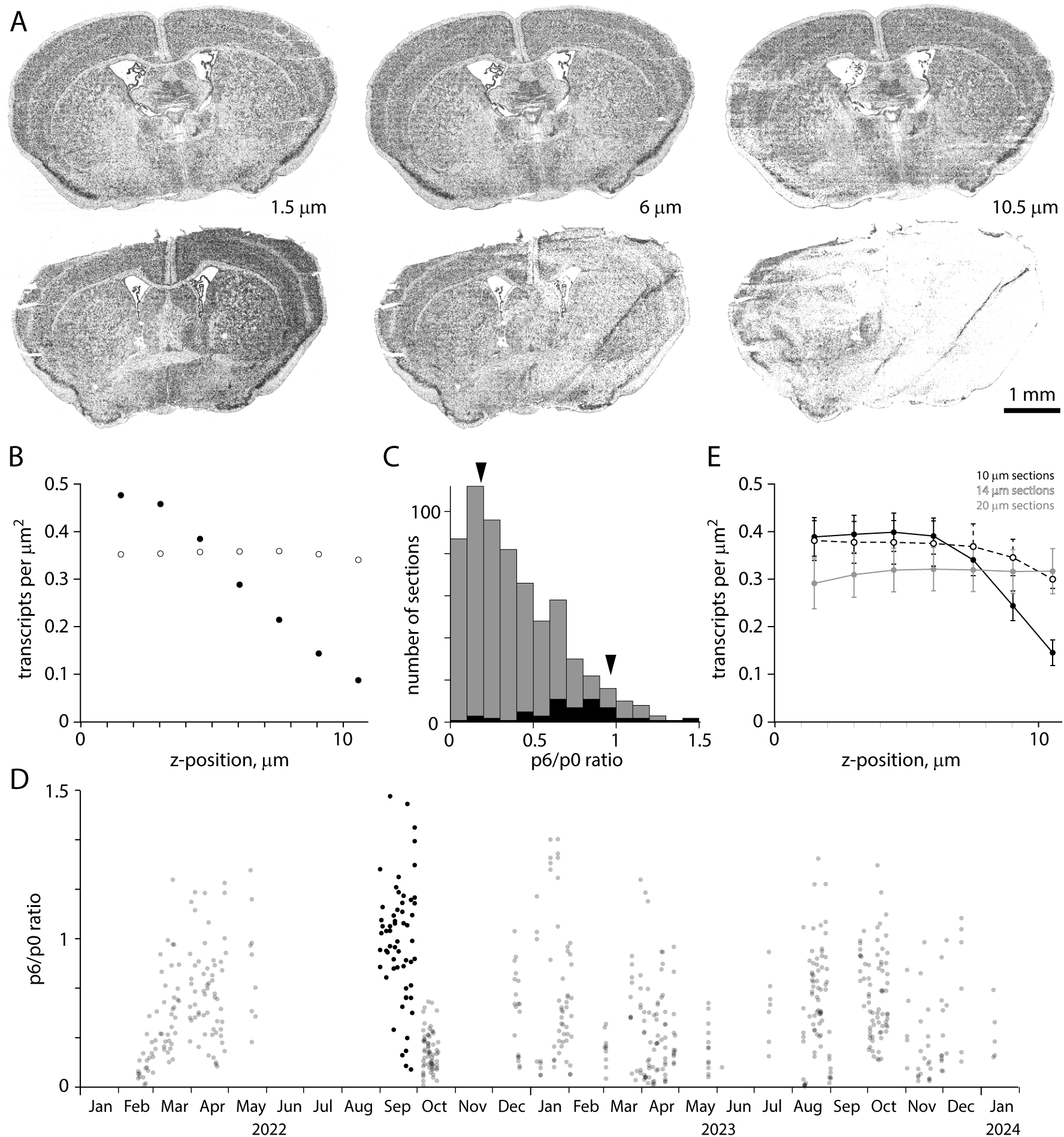
Uneven detection efficiency along the optical axis. (A) Transcript locations in three z planes for each of two neighboring sections from the same mouse brain, separated along the A-P axis by 200 µm. Distances are from the coverslip surface. (B) Transcript counts along the z-axis for the section in panel A. (C) Distribution of p6/p0 ratio for 641 sections (grey) and the 59 sections in the Yao et al. (2023) Allen Brain Cell Atlas dataset (black). Arrowheads, sections in A. (D) p6/p0 ratio over time. (E) Mean ± SEM transcript counts along the z-axis for 10, 14 and 20 µm thick sections. 3 sections each at 14 and 20 um, 6 at 10 um.

The p6/p0 ratio was skewed towards 0 with a median of 0.34 (mean of 0.39, figure 6C). The p6/p0 ratio distribution was shifted towards 1 for sections in the Yao *et al*. (2023) Allen Brain Cell Atlas dataset, with a median of 0.74. p6/p0 ratio changed little over time (figure 6D). To improve homogeneity in detection efficiency in the imaged volume, we cut thicker tissue sections while maintaining the number and separation of imaging planes (7 planes at 1.5 µm intervals, figure 6E). The p6/p0 ratio was ∼1 for 20 µm sections, but at a cost of fewer transcripts close to the coverslip and fewer total transcripts (mean ± SEM transcript count per µm^2^, summed along the z-axis: 2.31 ± 0.20 for six 10 µm sections, 2.53 ± 0.28 for three 14 µm sections, 2.20 ± 0.33 for three 20 µm sections). 14 µm thick sections proved a good compromise, with transcript numbers near the coverslip comparable to 10 µm sections (0.38 ± 0.04, 3 sections vs 0.39 ± 0.04, 6 sections) and a p6/p0 ratio of 0.79 ± 0.04 (three sections, vs 0.42 ± 0.11 for six 10 µm sections).

The decline in transcripts with distance from the coverslip differed between sections from a brain so tissue quality is unlikely to be a major factor. Our results provide little further insight into possible mechanisms, but the access of solutions to the deep (near the coverslip) and superficial faces of the section differ during benchtop chemistry and on the MERSCOPE, with the deep face being less accessible than the superficial face. Loss of RNA from the section, preferentially from the superficial face, might cause the gradient in transcript detection. Similarly, unbinding and loss of readout probes into wash solution during imaging would have a similar effect. Although the mechanism is unclear, detection of transcripts is rarely uniform through the depth of a MERSCOPE section.

In summary, the detection of transcripts in MERSCOPE experiments is rarely homogenous, varying in all 3 spatial dimensions and between sections. Our simulations provide some sense of the magnitude of the resulting effects on cell labels, but the variation in detection efficiency is complex enough that it’s likely not possible to map the accuracy of cell type labels throughout a section. More homogenous detection efficiency would facilitate the interpretation of spatial results.

Across our fleet of 8 MERSCOPEs, we observed significant differences in the magnitudes of all imperfections (ANOVA, p<0.05), but differences were slight. Overall, performance was similar across MERSCOPEs.

### Visual inspection

MerQuaCo characterizes the most common imperfections in each section, based on the transcript table. There are imperfections that are not detected by MerQuaCo, most often imperfections that are not evident in the transcript table or imperfections that become apparent when comparing nearby sections. For every experiment, we view results in the MERSCOPE Vizualizer, manually searching for imperfections. Figure 7 provides two examples of imperfections that were rare, not detected by MerQuaCo, but were observed multiple times by manual inspection. Figure 7A-C illustrate data loss in the DAPI image, visible as horizontal stripes through the left half of the section (figure 7A, B) and resulting in the local loss of somata within the image, and transcripts that cannot be assigned to a soma. Although DAPI information is lost, transcripts are observed throughout the section (figure 7C), preventing this imperfection from being detected by inspection of the transcript table or a transcript image.

**Figure 7.**
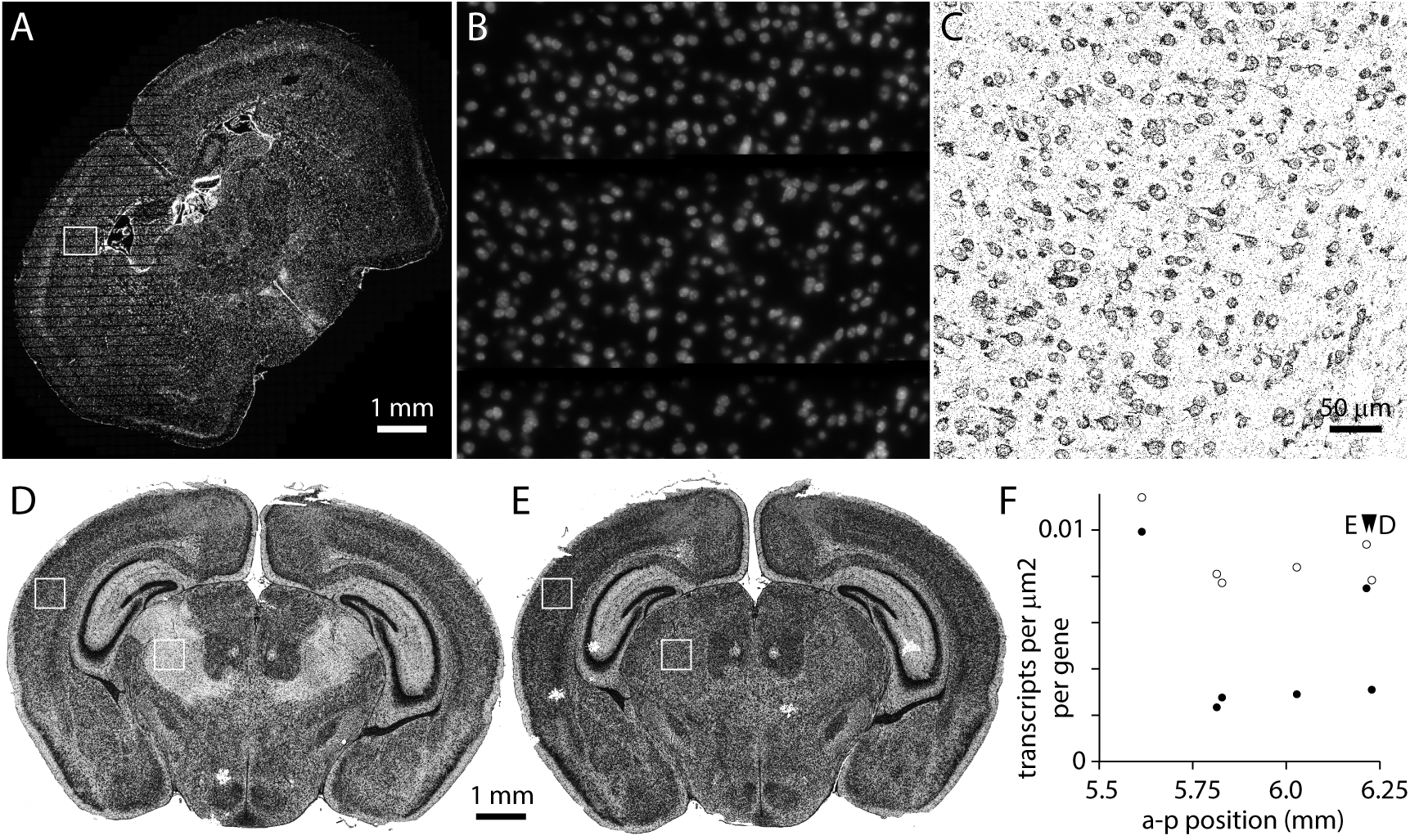
Imperfections identified by manual inspection. (A) DAPI image. Data loss results in horizontal stripes in the left half of the image. (B) DAPI in the sub-region in the box in panel A. (C) Transcripts in the corresponding region. (D) Transcripts in a section 6.2 mm posterior to bregma. (E) Transcripts from a neighboring section. (F) Transcript density in cortex and thalamus (boxes in panels D and E) for 6 neighboring sections.

**Figure 8.**
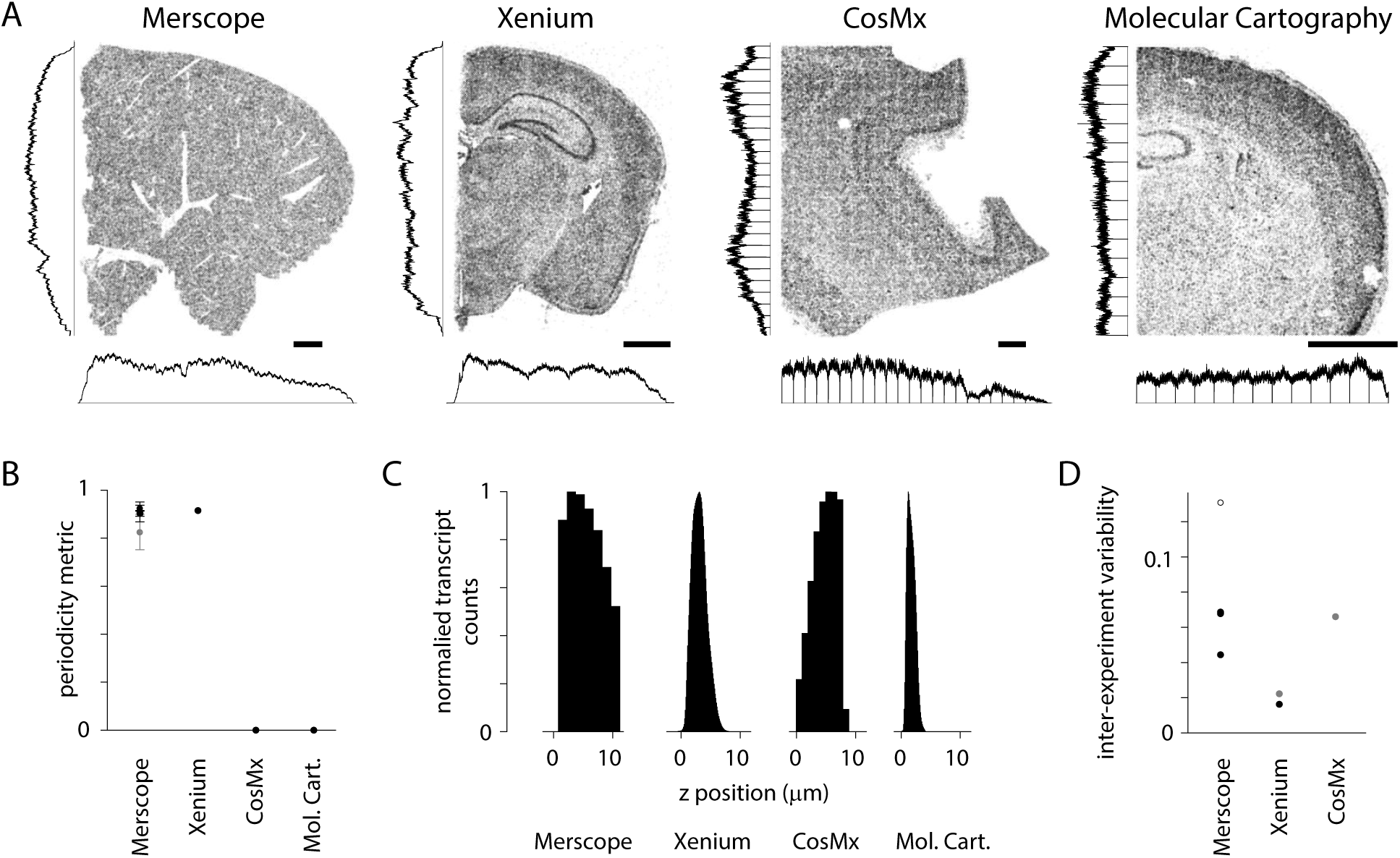
Transcript density across spatial transcriptomics platforms. (A) Example sections from four commercial platforms. MERSCOPE, mouse liver section. CosMx, human brain. Xenium and Molecular Cartography, mouse brain. Scale bars, 1 mm. Histograms indicate transcript density in (cardinal axes, normalized to peak). (B) Periodicity metric for public datasets (4 sections in A and 3 Vizgen mouse brain datasets). Grey: mean ± stdev periodicity metric for the Yao et al. (2023) Allen Brain Cell Atlas dataset. (C) Transcript count along the z axis for sections in A. (D) Pairwise CV of transcripts across experiments. MERSCOPE, 3 Vizgen mouse brain datasets and Yao et al. (2023) Allen Brain Cell Atlas dataset (open symbol). Xenium, fresh-frozen-mouse-brain-replicates-1-standard dataset from 10x. Grey datapoints (Xenium and CosMx) from Cook et al. (2023).

Occasionally we observed imperfections that are evident only when comparing sections. For example, in figure 7D-F a region of thalamus is missing transcripts in one section (figure 7D). An abrupt change in transcript density running along an anatomical boundary might result from localized expression, but in this instance the neighboring section displays no comparable change in transcript density (figure 7E). Moreover, transcripts are lost from thalamus in 4 of 6 neighboring sections (figure 7F). Clearly these differences are not biological: detection is reduced >50% in thalamus in 2 of 6 sections, likely resulting in a marked decline in accuracy with which cell populations in thalamus can be identified in these sections. MerQuaCo operates on individual sections so will not detect imperfections that are evident only when comparing sections. We search for intra-section changes in detection probability manually, by comparing results from nearby sections.

### Variations in transcript density on commercial spatial transcriptomics platforms

Many of the imperfections described above occur in datasets collected with multiple spatial transcriptomics platforms. We examined publicly accessible datasets from four commercial spatial transcriptomics platforms: Vizgen MERSCOPE, 10x Genomics Xenium, NanoString CosMx, and Resolve Molecular Cartography. For some sections, uneven detection across the field of view was visible by eye and was captured by our periodicity metric (figure 8A, B). For all sections, transcript density varied along the z axis (figure 8C). For platforms where results from multiple sections were available, we estimated differences in transcript count between sections, a proxy for inter-experiment differences in detection efficiency (figure 8D). With only small numbers of sections available, the results of this comparison should be considered preliminary but our results indicate that the imperfections we have described, and that we detect and quantify with MerQuaCo, occur on spatial platforms other than MERSCOPE. In some instances, imperfections are pronounced, underlining the potential value of applying MerQuaCo to other platforms.

## DISCUSSION

Here we have documented the incidence and magnitude of imperfections in image-based spatial transcriptomics datasets, focusing on the most common imperfections in a dataset of hundreds of sections collected over ∼2 years on the MERSCOPE platform. In time, these imperfections may be eliminated by equipment manufacturers, but not all the technical challenges have been solved in this new and rapidly evolving field and there is a need to characterize imperfections that persist in spatial datasets. Unfortunately, residual imperfections are often not obvious upon inspection of transcript or cell images. Our code, MerQuaCo, allows the user to detect and visualize imperfections, assisting in the process of quality control.

Like many other groups, we use gene expression profiles from spatial datasets as the basis for cell type labels (e.g. Yao *et al*., 2023). Most imperfections do not prevent the identification of cell types but impact the accuracy of labels, reducing confidence in labels and perhaps limiting the granularity with which cell populations can be characterized. Which imperfections have the greatest impact on the accuracy of cell type labels? Tissue damage and detachment from the coverslip, both of which result in local data loss, prevent all downstream analyses for the affected regions of the section. These two imperfections affect all transcripts and are therefore obvious on visual inspection of the dataset and are unlikely to lead to hidden errors in interpretation.

Like tissue damage and detachment, dropped images result in local data loss. In contrast with tissue damage and detachment, typically dropped images result in loss of only a subset of transcript species. This is a critical difference since many probe panels designed to identify cell types include some redundancy. In our experiments, dropped images eliminated tens of genes from a panel of 500. Often, the effect on the accuracy of cell type labels is modest, particularly for class and subclass labels. Dropped images may be more problematic where the aim of the experiment is other than cell typing. Where the aim is to measure the expression of one or a small number of genes, for example, dropped images may simply eliminate information on the genes of interest in affected regions.

When using spatial transcriptomics to locate genetically defined cell populations, the most impactful imperfections are differences in transcript density between sections and through space within a section. Our results indicate that transcript densities differ ∼2-fold between sections, ∼30% along the x and y axes, and ∼5-fold in z, and these effects are presumably multiplicative. The consequences can be substantial. Perhaps only ∼50-60% of cluster labels are accurate near the surface of a typical section. Furthermore, the effects of local changes in transcript density are difficult to assess. One solution might be to discard results from sections or from regions of a section where transcript density drops below a critical threshold. This threshold will depend on the aims of the experiment, but MerQuaCo could facilitate such a solution by quantifying transcript density.

Our analyses of publicly accessible datasets indicate that some of the most common imperfections in our MERSCOPE dataset also occur on other platforms. The accessible datasets are relatively small, often a few sections, sometimes just part of a section, preventing a thorough comparison of imperfections across platforms. As a result, our analyses only hint at the some of the relative strengths and weaknesses of different platforms. We expect that, as with MERSCOPE, imperfections will differ across experiments on each platform, necessitating quality control to identify experiments that meet the needs of the study. MerQuaCo could form the basis of such a quality control process, with only minor changes to the code needed to enable the analysis of results from other platforms.

Previous authors have compared results across spatial transcriptomics platforms, focusing on high-dimensional analysis of transcripts and cell expression profiles (Cook *et al*., 2023; Wang *et al*., 2023; Hartman & Satija, 2024). Cook *et al*. (2023) compared Xenium and CosMx results from prostate adenocarcinoma samples; Wang *et al*. (2024) compared MERSCOPE, Xenium, and CosMx results with FFPE tissue from multiple organs; and Hartman & Satija (2024) compared results from fresh-frozen mouse brain slices across 6 spatial transcriptomics platforms.

These authors focused primarily on platform-specific differences in transcript specificity and sensitivity, cell boundary identification, and the resulting differences in cell RNA content and classification. A consistent conclusion was that results were generally reproducible, across samples processed on each platform, and across platforms. These authors discussed the criteria by which they might select datasets for further analysis, and discard others, implying that there is enough variability between experiments that not all datasets are equally informative. For example, Wang *et al*. (2024) compared transcript counts per gene and pairwise correlation coefficients and suggested that these measures might form the basis for decisions on which datasets to include or discard. However, it remained unclear which parameters might best differentiate higher and lower quality datasets, and how to use these parameters to select the highest quality datasets.

Our results indicate that variability across experiments is substantial and the consequences for downstream interpretation can be significant, making it difficult to compare platforms. Firstly, one needs a large enough dataset from each platform to differentiate between results of differing quality since collecting a small number of samples from each platform leaves open the possibility that experiments were unusually successful on one platform and unusually unsuccessful on another and that the resulting comparison misleads. Secondly, one needs to develop and apply inclusion criteria for each platform that support a fair comparison. To better understand how spatial transcriptomics platforms compare, we need additional studies based on large datasets with intentional filtering of datasets before comparison. In short, our analyses complement previous studies, quantifying imperfections in spatial results rather than comparing platforms. Combining our approach and those described by previous authors would likely bring further insight into the relative performance of different platforms.

Despite the imperfections we’ve documented, numerous groups have published reliable results with spatial methods. An example is our recent atlas of cell types in the adult mouse brain (Yao *et al*., 2023). Many imperfections exist in our published dataset, documented here. In our experience, the presence of imperfections rarely prevents the collection of valuable results with spatial platforms. Rather, we regard the characterization of imperfections as a quality control step that allows the identification and perhaps elimination of the weakest datasets and the accurate interpretation of results, aware of the remaining imperfections and their possible consequences. Ideally our study of imperfections combined with future studies will build a consensus and more software tools for quality control, standardizing and streamlining data processing and further enhancing the reliability of results, thereby facilitating more discoveries with spatially resolved molecular imaging methods.

